# Denitrification in foraminifera has ancient origin and is complemented by associated bacteria

**DOI:** 10.1101/2021.12.27.474233

**Authors:** Christian Woehle, Alexandra-Sophie Roy, Nicolaas Glock, Jan Michels, Tanita Wein, Julia Weissenbach, Dennis Romero, Claas Hiebenthal, Stanislav N. Gorb, Joachim Schönfeld, Tal Dagan

## Abstract

Benthic foraminifera are unicellular eukaryotes that inhabit sediments of aquatic environments. Several foraminifera of the order Rotaliida are known to store and use nitrate for denitrification, a unique energy metabolism among eukaryotes. The rotaliid *Globobulimina* spp. has been shown to encode an incomplete denitrification pathway of bacterial origin. However, the prevalence of denitrification genes in foraminifera remains unknown and the missing denitrification pathway components are elusive. Analysing transcriptomes and metagenomes of ten foraminiferal species from the Peruvian oxygen minimum zone, we show that denitrification genes are highly conserved in foraminifera. We infer of the last common ancestor of denitrifying foraminifera, which enables us to predict the ability to denitrify for additional foraminiferal species. Additionally, an examination of the foraminiferal microbiota reveals evidence for a stable interaction with *Desulfobacteracea*, which harbour genes that complement the foraminiferal denitrification pathway. Our results provide evidence that foraminiferal denitrification is complemented by the foraminifera-associated microbiome. The interaction of foraminifera with their resident bacteria is at the basis of foraminiferal adaptation to anaerobic environments that manifested in ecological success within oxygen depleted habitats.

**Significance Statement:** A substantial component of the global nitrogen cycle is the production of biologically inaccessible dinitrogen attributed to anaerobic denitrification by prokaryotes. Recent evidence identified a eukaryote – foraminifera – as new key players in this ‘loss’ of bioavailable nitrogen. The evolution of denitrification in eukaryotes constitutes a rare event and the genetic mechanisms of the denitrification pathway in foraminifera are just starting to be elucidated. We present large-scale sequencing analyses of ten denitrifying foraminiferal species, which reveals the high conservation of the foraminiferal denitrification pathway. We further find evidence for a complementation of denitrification by the foraminiferal microbiome. Together, these findings provide novel insights on the early evolution of a previously overlooked component in the marine nitrogen cycle.

## Introduction

Nitrogen is an essential element for life on earth as it forms the basis for the synthesis of nucleotides and amino acids. Nonetheless, while the earth atmosphere is rich in nitrogen gas (up to 78%) (1), this gas is usually inert but can be made accessible for biological processes by nitrogen fixation (2). Microbial organisms are important players in the global nitrogen cycle as they facilitate the assimilation of nitrogen into bioavailable nitrogen derivatives as well as the dissimilation of nitrogen derivatives into dinitrogen (N _2_) (2). A key dissimilatory pathway is denitrification, where nitrate (NO _3-_) is either partially or completely degraded and the final product – N _2_ – is released to the atmosphere (i.e., nitrogen loss) (2). Marine organisms are considered major contributors to nitrogen loss from the environment with benthic organisms being responsible for about two-thirds of the loss of reactive nitrogen in the ocean (3, 4). Especially oxygen minimum zones (OMZs) are worth mentioning here as they are estimated to be responsible for 20-40% of bioavailable nitrogen removal in the ocean (3, 5). The ability to perform denitrification is abundantly found in eubacteria (6), whereas it is rare among the eukaryotes. Partial or complete denitrification have only been reported for two species of fungi (7) and several foraminifera of the order Rotaliida (8-11). Denitrifying foraminifera are unicellular eukaryotes commonly found in marine sediments. Studies of foraminifera residing in the Peruvian OMZ showed that they are found in high densities of up to 600 individuals per cm^2^, where they are estimated to contribute 20%-50% to the total benthic nitrate loss in the OMZ (10, 12).

Rotaliid foraminifera are traditionally divided into three clades based on their phylogeny (13). Only species in clades I and III were demonstrated to denitrify, while rotaliids classified in clade II have been shown to lack an intracellular nitrate storage or measurable denitrification activity (9). Class II rotaliids typically populate environments with poor-nitrate supply, e.g. intertidal to near-shore habitats from the tropics up to boreal bioprovinces (14), hence it is conceivable that they lack the ability to denitrify. One example is *Ammonia tepida* (class II), that can survive episodic oxygen depletion events via dormancy (15). A study of clade III species sampled from a hypoxic environment (Gulmarfjord, Sweden) – *Globobulimina* spp. – showed that that their genome encodes several genes along the denitrification pathway that are of ancient bacterial origin (11). These include the copper-containing nitrite reductase (NirK) and nitric oxide reductase (Nor), but not the nitrate reductase (NapA/NarG) and nitrous oxide reductase (NosZ). In addition, the *Globobulimina* genome contains a diverse gene family encoding for nitrate transporters (Nrt) (11), a finding that is consistent with the accumulation of an intracellular NO _3-_ storage in denitrifying rotaliids (8-10, 16). More recent studies reported the presence of those genes in at least two additional species of foraminifera and provide alternative suggestions on the nitrate reductase homolog missing from the earlier model of denitrification in foraminifera (17, 18). The rotaliids ability to respire both oxygen (O _2_) and NO _3-_ marks them as facultative anaerobes that are able to thrive in both aerobic and anaerobic conditions. Furthermore, while oxygen is generally considered to be preferred over NO _3-_ as electron acceptor, rotaliids from the Peruvian OMZ were reported to prefer NO _3-_ over O _2_ (10). These findings suggest that genes of the denitrification pathway are widespread in rotaliids, however their distribution remains largely unknown.

Foraminifera, like other eukaryotes (19), are populated by bacterial organisms that reside out-and in-side their cell (i.e., test) (20-24). Microscopic observations showed that the bacteria are localized in food vacuoles or in the cytoplasm of the cell, which led to the suggestion that interactions of foraminifera with its microbiota may vary from prey-predator interactions or parasitism to metabolic symbiosis (22). Further studies of the microbiota in clade II rotaliids identified sulphate-reducing and sulphur-oxidizing bacteria that were suggested to participate in Sulphur cycling thus providing carbon and other nutrients to the host (20, 21, 25). Notwithstanding, previous studies of foraminifera-associated bacteria used microscopy observations or sequencing of marker genes (e.g., 16S rRNA) that are limited in their resolution (20-25). It has been suggested that the microbiota of denitrifying foraminifera include denitrifying bacteria that utilize the foraminiferal NO _3-_ storage, similarly to observations in other bacteria-protist associations [e.g., gromiids (26), allogromiids (27)]. Indeed, metagenomics of the *Globobulimina* microbiota revealed the presence of a taxonomically diverse species community where several members encode homologs of NapA and NosZ (11). This finding gave rise to the hypothesis that the partial foraminiferal denitrification pathway may be complemented by microbiota functions.

Here we investigate the evolutionary history of genes along the denitrification pathway in foraminifera and furthermore examine the functional repertoire of the foraminiferal microbiome. For that purpose, we studied populations of ten rotaliid species known to denitrify (10) (except *Globobulimina pacifica* for which no information is available). Our study supplies insights in the evolution of rotaliids and their microbiota and lays the basis for further research of foraminiferal genome evolution.

## Results

### Transcriptomes of ten Peruvian rotaliids

For the purpose of our study, we sampled benthic foraminifera in the Peruvian oxygen minimum zone. Individual foraminifers from among ten focal species were manually identified and picked using a stereomicroscope (see sampled species in Figure 1A). Eukaryotic transcriptomes and whole metagenomes were sequenced from the sampled foraminifera with two biological replicates per species (ca. 160 individuals per sample). Our computational analysis includes five additional publicly available transcriptomes of foraminifera (Suppl. Table S1) and the genome data of one monothalamid species (*Reticulomyxa filosa*). The assessment of transcriptome completeness showed that ∼90% of the eukaryotic marker proteins are present in the data of the ten newly sequenced species. Furthermore, we examined the purity of the focal species in each transcriptome by testing for redundancy of eukaryotic marker proteins that are expected as single-copy genes. As single-copy genes are expected to exist only once in a given species, their redundancy might indicate for additional eukaryotic species present in our samples. The highest proportion of such bystander species were found in the *B. spissa* (78%) and *B. costata* (50%) transcriptomes (Suppl. Table S1).

**Figure 1.**
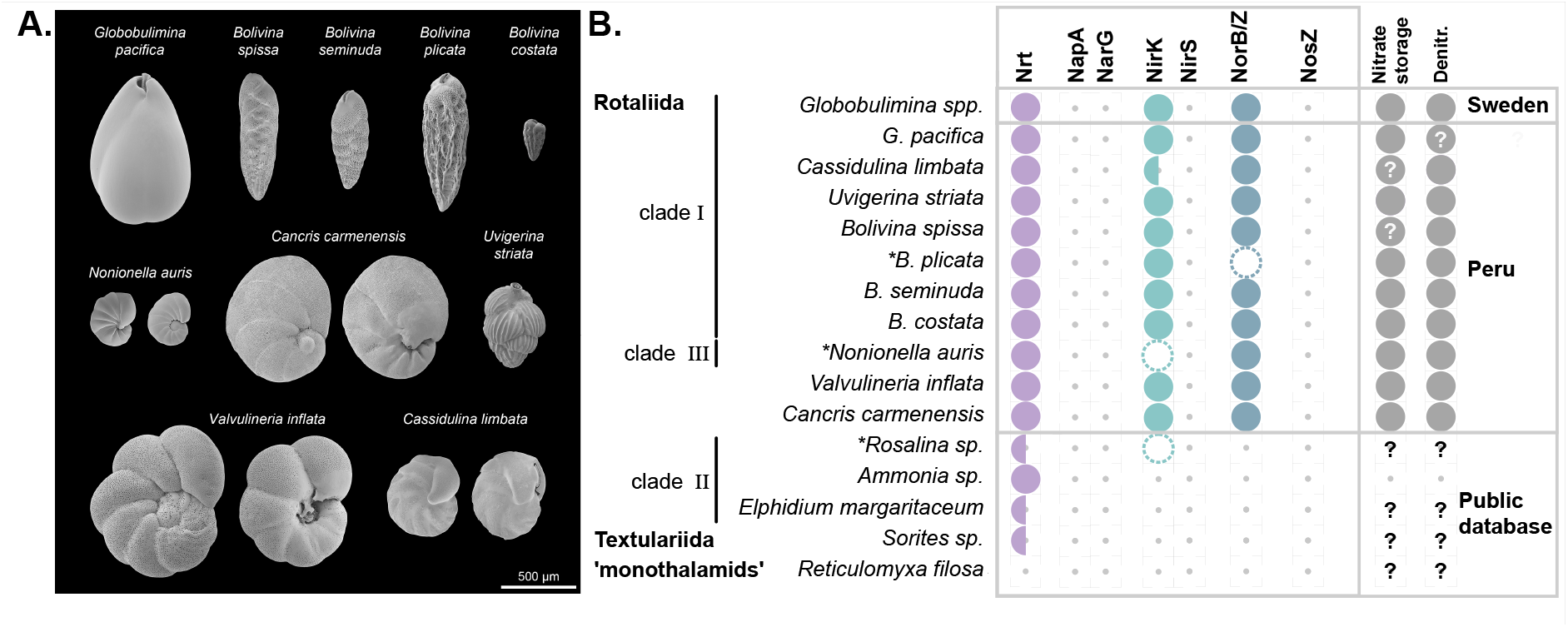
Morphological characteristics of the sampled rotaliids and denitrification gene repertoire. **A)** Scanning electron micrographs of the sampled species. Species with clearly distinct lateral views are shown from two sides. For more detailed views see Suppl. Figure S1. **B)** Presence/absence of homologs of denitrification proteins identified for different foraminifera. Half circles indicate species where only a single, of multiple subtypes, was found. Dashed circles illustrate homologs discarded due to low coverage and asterisks highlight corresponding species. Evidence for NO3^-^storage and/or denitrification activity is illustrated by closed circles in the last column. Question marks denote missing information. Foraminifera sampled in this study are highlighted by the location ‘Peru’. Protein symbols: Nrt: nitrate/nitrite transporter; NapA: periplasmic nitrate reductase; NarG: membrane-bound nitrate reductase; NirK: copper-containing nitrite reductase; NirS: cd1-containing nitrite reductase; Nor: nitric oxide reductase; NosZ: nitrous oxide reductase. Note that the identification of homologous genes is based not only on sequence similarity but also on transcript abundance in order to exclude bystander species in the data. The additional data filtration stage affected our findings for *Rosalina* sp., *B. plicata* and *N. auris* (Suppl. Table S2 & S3), where the presence of at least one the crucial homologs (i.e, NirK or Nor) remained in the *B. plicata* and *N. auris* metatranscriptomes.

### The Peruvian rotaliids harbour genes of the denitrification pathway

To explore the denitrification mechanisms in the Peruvian foraminifera, we searched for homologs to denitrification pathway genes. Our results reveal that all sampled species harbor homologous genes to Nrt, NirK and Nor (Figure 1B; note that genes are termed by their corresponding protein symbol). Our search for homologs in the publicly available data yielded a putative homolog of NirK in *Rosalina* sp. and Nrt homologs in all but *R. filosa* (Fig. 1B). The additional species we included here, especially *Ammonia sp*., *Elphidium margaritaceum* and *Sorites sp*., are considered to reside mostly in oxygenated habitats, hence they are not expected to be able to perform denitrification. These species lack homologs to the denitrification genes, but encode nitrate transporters (Nrt, Figure 1B). Among this group, only *A. tepida* was so far experimentally tested for denitrification ability and reported as a non-denitrifying species (9). The reconstructed phylogenies indicate that all the examined genes are indeed encoded in the foraminiferal genomes, as all of them group with homologs previously reported to originate from the nuclear genomes of *Globobulimina* spp. (11). Furthermore, the phylogenies of Nrt and NirK, reveal two major sub-clades including most, but not all, of the species sampled (Full and half circles in Fig. 1B; Suppl. Figure S2; Suppl. Table S2). Previously we reported the presence of subclades in the Nrt and NirK phylogenies (11), but could not clarify if those were explained by sequencing errors, gene duplication, or speciation events. The phylogenetic trees presented here show that each subclade is represented in most of the rotaliids sampled in the current study. Consequently, we conclude that ancient gene duplications in the Nrt and NirK gene families led to the evolution of two or more sub-clades that encode for different (paralogous) protein subtypes. Overall, our results demonstrate that the denitrification pathway is highly conserved in the Peruvian rotaliids.

### Foraminiferal denitrification evolved in the Rotaliida ancestor

The presence of NirK and Nor homologs the tested rotaliids suggests that those genes may have an ancient origin in this order. To study the origin of denitrification in foraminifera, we reconstructed the phylogenetic relations among species in our data using 81 eukaryotic marker proteins that have homologs in all species analyzed here (Suppl. Figure S3; Suppl. Table S4). Here, we applied a phylogenomics approach where the root position is inferred from all marker genes independent of a single species tree (28) (see methods). Our results show that the best-supported root position (42% of the gene trees) was at the branch leading to *R. filosa* (Suppl. Figure S3). This result is in agreement with previous studies that assumed monothalamids to be an outgroup in foraminiferal phylogenies (29). The second most frequent root position was found at the branch leading to the miliolid *Sorites* sp. (15% of single gene trees), which was followed closely by a root position on the branch leading to *Rosalina* sp. (12%). Taken together, the inferences of rooted topologies show that the Peruvian rotaliids, together with *Globobulimina* spp. form a monophyletic group. Thus, our results reveal a shared origin of denitrification in foraminifera and hence the evolution of denitrifying species within the order Rotaliida.

For the inference of the denitrifying rotaliids last common ancestor (LCA; i.e., the rotaliids root), we reconstructed phylogenetic trees from an extended set of 146 eukaryotic marker protein coding genes considering only members of the Rotaliida (Figure 2; Suppl. Table S4). An examination of the rooted tree topologies revealed that the branch leading to *Ammonia* sp. and *E. margaritaceum* was the most frequently supported root position (14% of single gene trees). This root inference is further supported by an alternative root position at a branch that splits *Ammonia* sp., *E. margaritaceum* and *Rosalina* sp. from the remaining species (10%). Overall, the different alternative root positions support with 35% a Rotaliida LCA within clade II foraminifera (i.e., among *Ammonia sp*., *E. margaritaceum & Rosalina* sp.; Fig. 2). The absence of denitrification genes in clade II members (Fig. 1) further supports the suggestion that the evolution of denitrification in rotaliids occurred by lateral gene transfer from a prokaryotic donor rather than being inherited from the rotaliid LCA (11).

**Figure 2.**
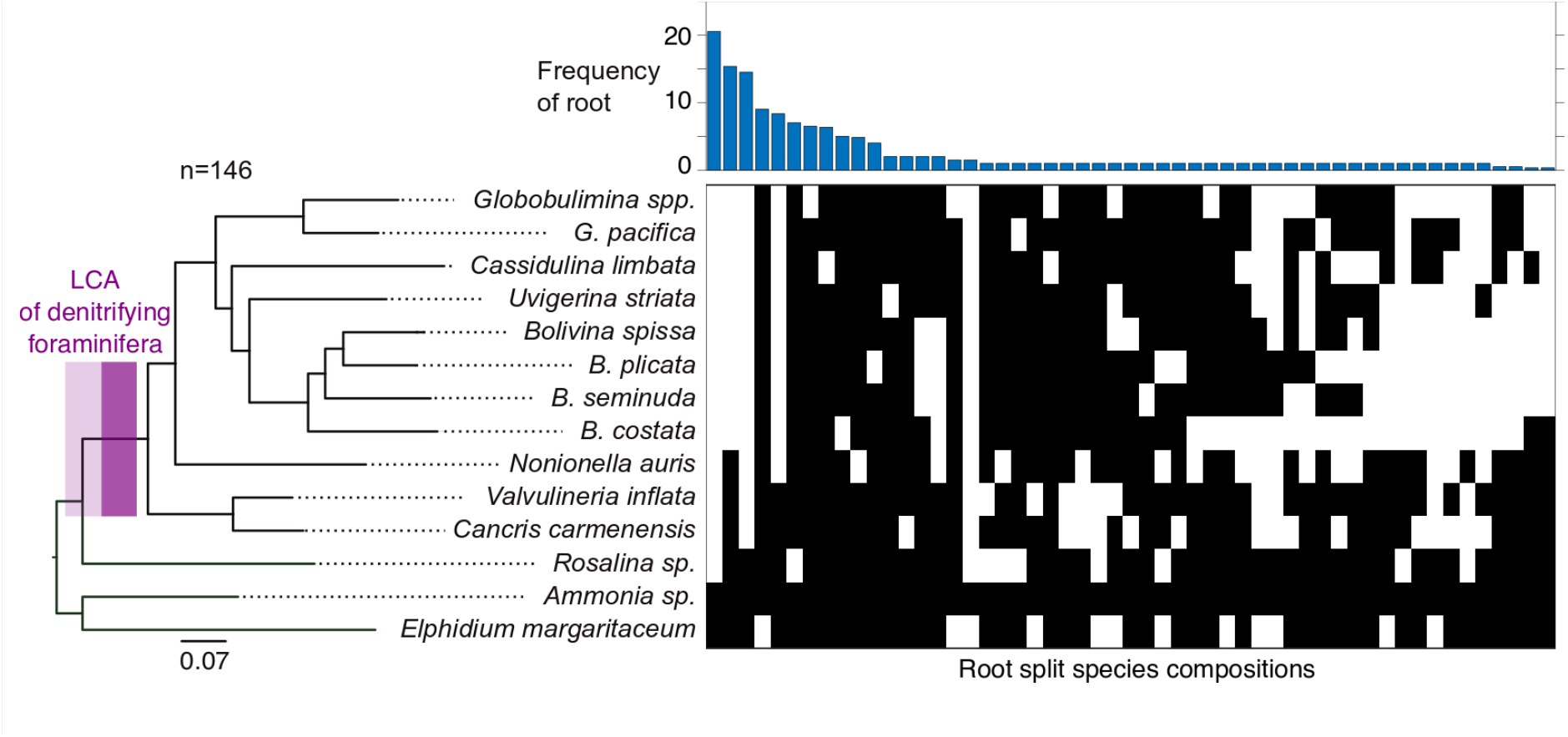
Phylogenetic rooting of rotaliids. A Maximum-likelihood phylogenetic reconstruction of foraminiferal species from 146 eukaryotic protein marker sequences is shown on the left. The branch with the LCA of denitrifying foraminifera is highlighted in purple, where a lighter color includes lower bound for the origin of denitrification. Nonparametric bootstrap support is 1000/1000 at all the branches. A split representation of root splits determined via the MAD approach for the 146 single gene trees is shown on the right. Each column represents a putative root branch reported as split of two groups (black and white boxes) indicating for the species found on either side of the branch. The bar graph reports the single gene tree count supporting the corresponding root split.

### Ancient origin of denitrification in foraminifera

To further study the denitrifying foraminifera LCA, we examined the phylogenetic position of the Peruvian species in the context of a foraminiferal species tree. For that purpose, we extracted sequences of the 18S ribosomal RNA (rRNA) subunit from our transcriptome data and used it in combination with publicly available 18S rRNA sequences to reconstruct a foraminiferal species phylogeny (Figure 3A; Suppl. Table S5). The resulting rooted topology was mostly in agreement with the rotaliids phylogeny, where the focal species are found next to their closest relatives. Most species grouped well with their previously defined clades with the exception of *Nonionella* and *Globobulimina* representatives. Furthermore, several taxa were not monophyletic, as has been previously observed in foraminiferal 18S rRNA phylogenies (29, 30).

**Figure 3.**
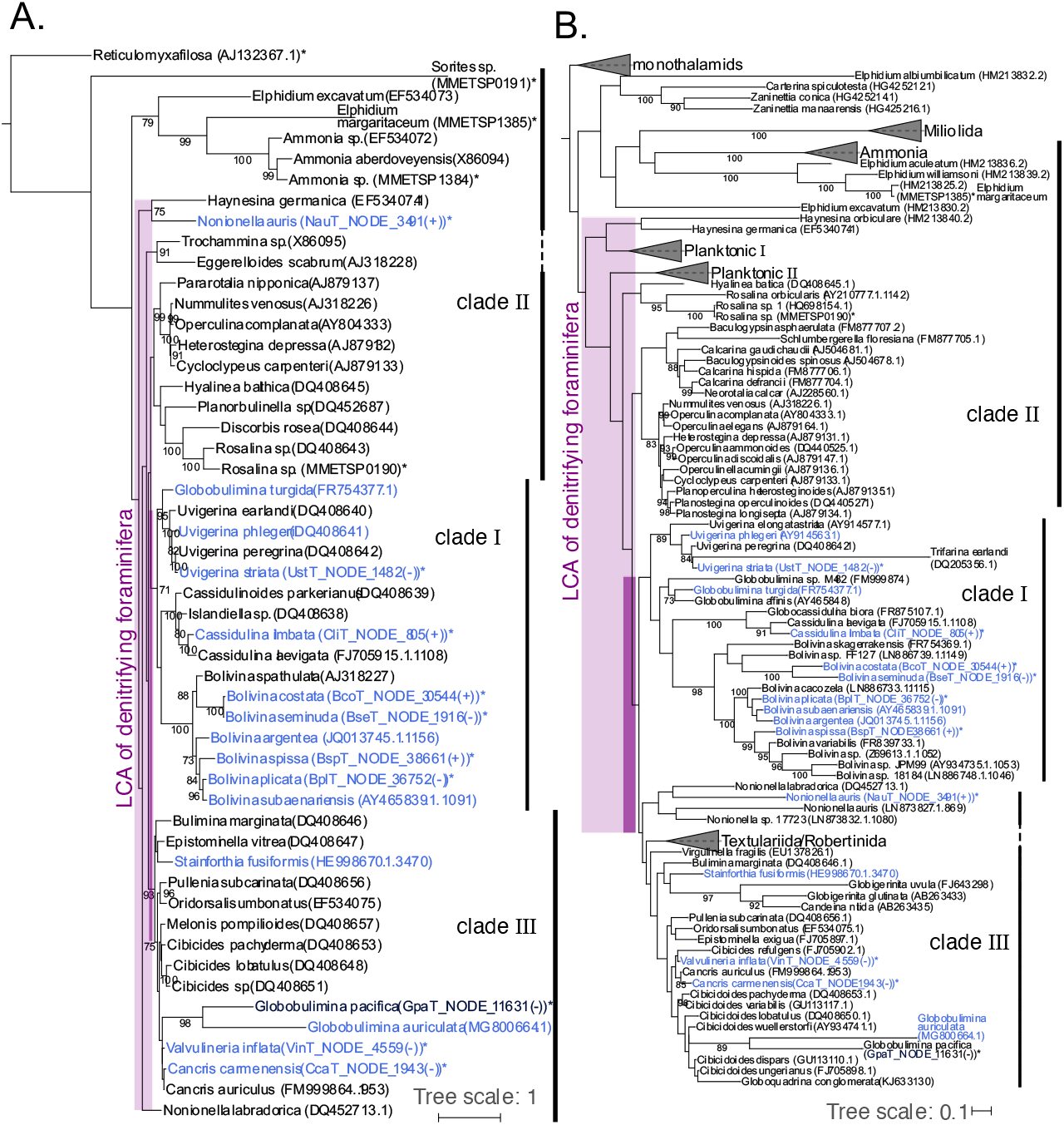
The origin of denitrification in context of the whole foraminiferal group. **A)** Large-scale phylogenetic representation of Peruvian foraminifera in context of different foraminiferal taxa based on 18S sequences available in public databases. Clades I & of the order Rotaliida appear are highly supported bootstrap values. Questionable branching of *Nonionella* outside class is not characteristic to data of the current study as *labradorica* that as well does not group directly with clad*Globobulimina pacifica* forms a clade with *G. auriculata*, but within clade However, we consider the branching of these two species as uncertain due to the long branches and the low bootstrap support, whichwe did not observe for the marker protein phylogeny (Figure 2). Lower (light purple; including *N. labradorica*) and higher (dark purple) boundaries for the origin of foraminiferal denitrification are highlighted by boxes. **B)** Extended phylogenetic representation of foraminifera including planktonic species. Lower (light purple) and higher (dark purple) boundaries for the origin of foraminiferal denitrification are highlighted by boxes. Planktonic I & III designate two to three distinct clusters comprising planktonic foraminifera. Species experimentally shown to denitrify are highlighted in blue. The only exception is *Stainforthia fusiformis*, where denitrification activity has been shown for an unspecified species of the same genus. Species also considered in Figure 1 or Suppl. Figure S3 are highlighted by asterisks. Several species labels contain as well the contig id and orientation of 18S sequences in the corresponding transcriptome assemblies. Bootstrap support values (>=70) with 1000 replicates are shown at the branches. The trees were rooted by the clade of monothalamids containing *R. filosa*. A detailed phylogeny is presented in Supplementary Figure S4.

The inference of the denitrifying foraminiferal LCA enabled us to further reconstruct the origin of denitrification within the foraminiferal species tree. For that purpose, we examined an extended phylogeny with additional foraminiferal groups including planktonic species (Figure 3B; Suppl. Figure S4). While the general taxonomic relationships in the tree were recovered as before (i.e., Figure 3A), many branches had a low statistical support. For example, the rotaliid clades II and III were paraphyletic, in contrast to the phylogeny with less taxa (Figure 3A). Nonetheless, an LCA of clades I and III could be inferred; the previously unassigned genera *Valvulineria* and *Cancris* (9) grouped well with members of clade III in both 18S rRNA phylogenies, hence they can be classified as clade III.

Since both clades I and III harbour several denitrifying foraminifera, it is most parsimonious to conclude that LCA of clades I and III was likely a denitrifying organism. Our inference thus suggests the presence of denitrification genes (and hence denitrification capability) in all members of clades I and III including e.g., the genera *Cibicoides* and *Virgulinella*, which were previously not considered as denitrifying species. Indeed, *Cibicidoides wullerstoerfi* has been assumed for a long time to be unable to withstand O _2_ depletion (31). However, recent studies observed living *Cibicidoides spp*. thriving in environments of < 2 µmol/kg O _2_ (32) and fossil specimens in the paleorecord during periods of severe O _2_ depletion (33). These recent studies support our prediction that several *Cibicidoides spp*. are able to denitrify.

### Composition of the foraminiferal microbiome is species-specific

To test for bacterial contribution to the foraminiferal denitrification, we examined the foraminiferal microbiome. For that purpose, we compared the composition of bacterial communities among foraminifera in our sample. In the absence of species-specific interactions, we would expect a strong impact of the environment (i.e., sampling location and water depth) on the microbiome composition. Thus, we compared the microbial community composition between foraminiferal species and their sampling depths (Figure 4A). Our results show that the individual microbiomes are clustered by the foraminiferal species rather than the sampling location, indicating the presence of species-specific bacterial communities in foraminifera. Our results reveal a similar taxonomic composition of bacterial communities associated with *Globobulimina* spp. from Sweden and *G. pacifica* from the Peruvian OMZ (Figure 4A). The similarity in microbiome composition among the *Globobulimina* species indicates the presence of genus-specific microbiome.

**Figure 4.**
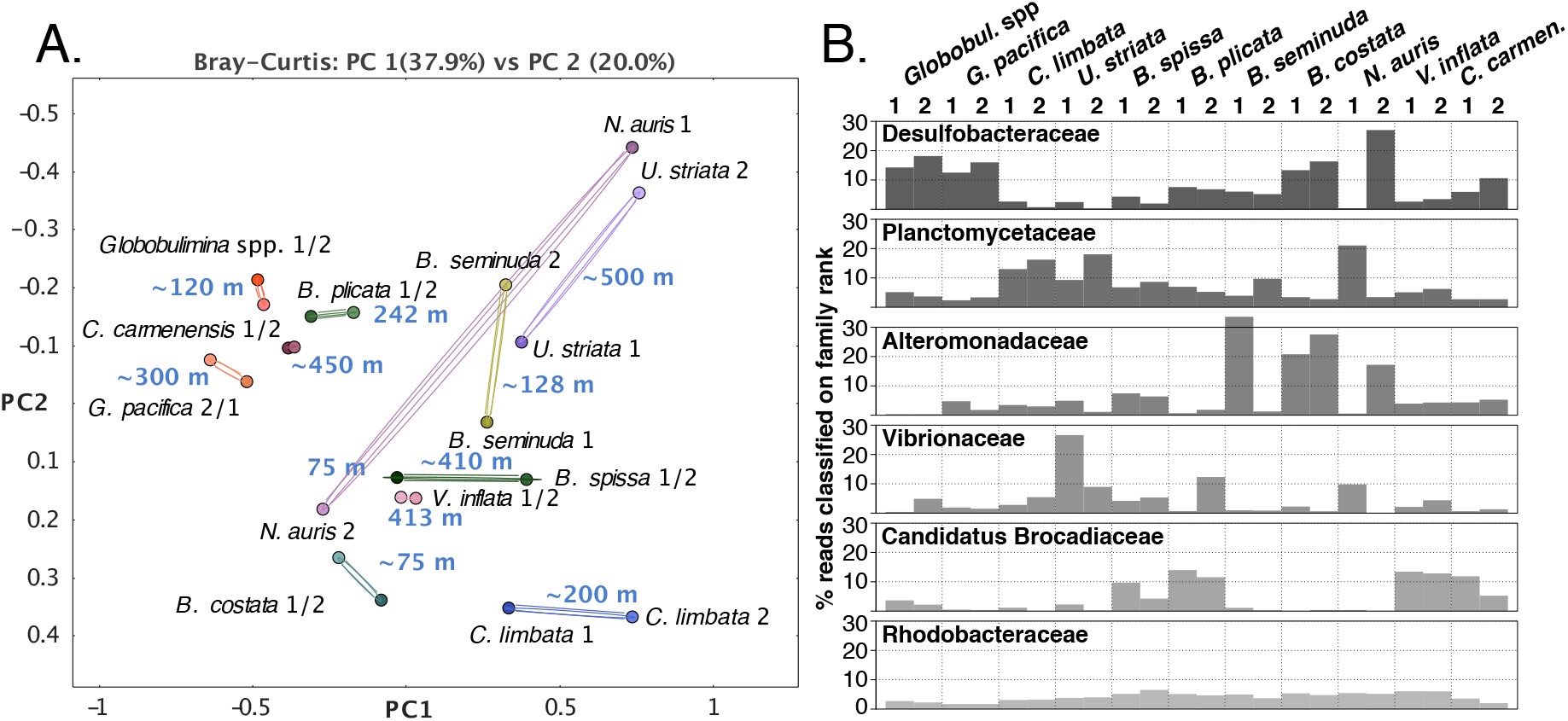
Comparison of microbial communities associated with different foraminifera. **A)** Principal coordinates analysis (PCoA) clustering of bacterial communities. Each circle represents a bacterial community; duplicates per species are connected by lines. The text next to the circles indicates for species name, replicate number or sampling depth. The clustering of most species-associated communities indicates for species-specificity. Grouping by higher taxonomy was only observed for *Globobulimina* spp. from Sweden and *Globobulimina pacifica* from the Peruvian oxygen minimum zone (OMZ). **B)** Bar graphs of the six most abundant bacterial families found in the foraminiferal bacterial communities beginning at the top. Each bar represents the proportion of reads assigned to the individual taxon relative to the total number of reads in a sample classified on family rank or lower. Note the prevalence of Desulfobacteraceae shared between the Globobulimina samples, while none of the other bacterial families showed high prevalence in the genus.

To identify common key players in foraminiferal microbiota we examined the relative abundance of all bacterial families in their microbiome (Figure 4B; Suppl. Table S6). Our results revealed that the most prevalent bacterial families are Desulfobacteracea followed by Planctomycetaceae. The relative abundance of these two families varied between different species and samples. Furthermore, communities with a high relative abundance of Desulfobacteracea are characterized by a low abundance of Planctomycteaceae and vice versa (*r* _*s*_=-0.77, p-value < 0.01, using Spearman correlation coefficient and t-test). Desulfobacteracea comprise sulphate-reducing bacteria and several representatives that are able to grow chemoautotrophically (34). Members of this group were previously observed in association with other rotaliids like *Virgulinella fragilis* and members of clade II (20, 25). Planctomycetes are often found in association with macroalgae (35); they are characterized by a diversified metabolism allowing them the of colonization of a wide range of habitats (35).

The next most abundant families were Alteromonadaceae, Vibrionaceae and ‘Candidatus Brocadiaceae’ that were more diverse in their relative abundances across different species and replicates. Most reported ‘Candidatus Brocadiaceae’ members are autotrophic, obligately anaerobic bacteria performing anammox, which is a dissimilatory pathway where ammonium and nitrite (NO _2-_) are metabolized into N _2_ (36). The presence of nitrate storage in foraminifera and NirK suggest that nitrite is readily available inside the foraminiferal test. Vibrionaceae are a diverse group including multiple species that colonize marine organisms either as symbionts (37, 38) or pathogens (39). Alteromonadaceae were isolated from diverse marine environments including eukaryotic microbiota (40). They are considered to be aerobe or facultative anaerobic bacteria that generally lack denitrification capabilities (41). Finally, another abundant family, Rhodobacteracea, is worth mentioning, as it is always found in low, but similar abundance in all samples. Marine Rhodobacteracea species are considered ecological generalists (42-44) and have been reported to colonize marine animals (e.g., fish larvae or sponges) (45, 46). The uniform distribution of Rhodobacteracea in the foraminifera-microbiome suggests that members of this group are permanent residents in the foraminiferal microbiota.

Our data thus shows that rotaliids are habitat to bacterial communities whose composition is akin to the microbiota of other marine eukaryotes.

### An ancient interaction between Desulfobacteraceae and *Globobulimina* hints to metabolic dependency

The metagenomes analysis and classification into bacterial metagenome-assembled genomes (MAGs) resulted in a total of 263 high-quality MAGs (i.e., draft bacterial genomes) of foraminifera-associated bacteria (Suppl. Table S1). The strongest signal for a stable core microbiome in our data was observed in the comparison among the *Globobulimina* species, whose microbiome is characterized by a high frequency of Desulfobacteraceae (Figure 4B). To further explore the association between *Globobulimina* and Desulfobacteraceae we examined the Desulfobacteraceae MAGs in our data. A total of 40 high quality MAGs were obtained from the *G. pacifica* metagenome, including four high quality draft MAGs classified as Desulfobacteracea. These were compared with the 26 previously published MAGs for *Globobulimina* from Sweden, which include two Desulfobacteracea MAGs (11) (Suppl. Table S7). A phylogenetic network of the Desulfobacteracea MAGs reveals two MAGs from the *Globobulimina spp*. and *G. pacifica* metagenomes (Glo_11 & Gpa_30) that appear as sister taxa (Figure 5A). The average nucleotide identity (ANI) between both MAGs was 84%, which is within the range expected for inter-species sequence similarity (e.g., within genera) (47). The common ancestry of these two MAGs suggests that the association between Desulfobacteracea and *Globobulimina* has an ancient origin. Our results thus indicate that the interaction fidelity between *Globobulimina* and Desulfobacteracea is high, similarly to observations in other eukaryote-bacterial symbioses [e.g., oligochaete worms and sulfur bacteria (48) or marine sponges with bacteria of the Poribacteria phylum (49)].

**Figure 5.**
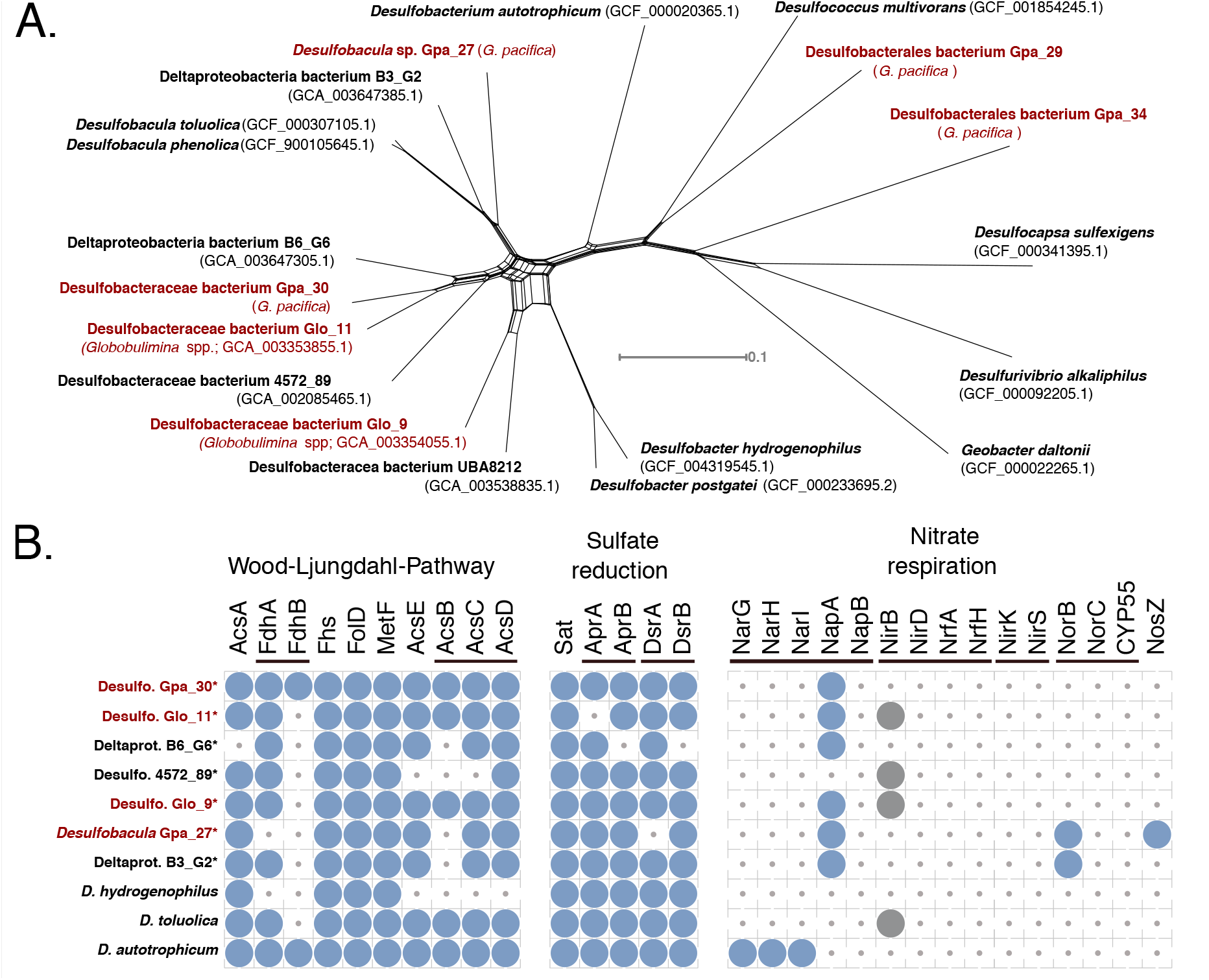
Characteristics of Desulfobacteraceae bacteria associated with Globobulimina. **A)** Neighbor-net network for Desulfobacteraceae associated with the genus *Globobulimina* from Sweden and Peru (*Globob-ulimina* spp. and *G. pacifica*, respectively) and additional Deltaproteobacteria. The network is based on a concatenated alignment of 33 single copy orthologous protein sequences found in all isolates shown. Red fonts highlight isolates found in association with members of the genus *Globobulimina*. The monophyly of the MAG Gpa_30 and MAG Glo_11 suggests a common ancestry. Note that the closest relatives to that pair are two uncultivated MAGs (Deltaproteobacteria bacterium B6_G6 & Desulfobacteraceae bacterium 4572_89) sampled from sediments in deep-sea hydrothermal vent sites at the Gulf of California. **B)** Presence of predicted proteins encoding metabolic pathways in representative members of Desulfobacteraceae. Three pathways are shown: Wood-Ljungdahl pathway, (dissimilatory) sulphate reduction & nitrate respiration. Asterisks highlight genomes in draft state. Circles indicate for the presence of homologs identified by the KEGG KAAS server. The thin bars link enzymes that participate in the same reaction step either by being an alternative enzyme or being part of one protein complex. The grey circles indicate for cases that were discarded following a manual inspection.

A central foundation in the evolution of symbiotic interaction is an exchange of currencies (i.e., resources) between the partners (50). The versatile metabolic capabilities of Desulfobacteraceae and the availability of intracellular NO _3-_ storage in foraminifera may suggest that the interaction between Desulfobacteracea and *Globobulimina* is based on nutritional currencies. To further examine possible symbiotic interaction between Desulfobacteracea and *Globobulimina*, we searched for metabolic properties of the Desulfobacteraceae MAGs that could serve as nutritional currencies in the symbiosis. For that purpose, we surveyed the metabolic pathways of representative Desulfobacteracea MAGs with respect to carbon, sulfur and nitrogen metabolism (Figure 5B; Suppl. Table S8). Our results reveal that all MAGs sampled encode the genetic repertoire required in order to perform dissimilatory sulfate reduction as well as the Wood-Ljungdahl pathway. Anaerobic respiration via the dissimilatory sulfate reduction to sulfide is widespread among Desulfobacteraceae. Members of this family that encode the Wood-Ljungdahl pathway are able to oxidize organic compounds to carbon dioxide (51). Alternatively, the Wood-Ljungdahl pathway may function in CO _2_ carbon fixation during autotrophic growth.

Genes along the denitrification pathway (or NO _3-_ respiration) are mostly absent from all Desulfobacteracea MAGs, except for periplasmic NO _3-_ reductases (NapA) homologs that were found in most foraminifera-associated MAGs (for a review in NapA function see ref. 52). All MAGs lack the typical NapB protein forming a complex with NapA. However, *nap* operons without NapB genes have been reported for other members of Deltaproteobacteria, e.g. *Desulfovibrio desulfuricans* (53). Notably, we found that the genome of *Desulfobulbus propionicus* (Accession: GCA_000186885.1), a member of the order Desulfobacterales that has been demonstrated to grow based on NO _3-_ (54), is lacking a gene for NapB. Hence NO _3-_ respiration in Desulfobacterales in the absence of NapB is possible. We found that the NapA sequences of all MAGs (and *D. desulfuricans*) include a signal peptide of the twin-arginine translocation pathway (reviewed in ref. 55) indicating that those proteins could be translocated across cellular membranes. It is tenable to hypothesize that NapA is secreted by the bacteria to *Globobulimina* intra-cellular environment. An alternative scenario is nitrate uptake by the bacterium followed by the reduction to nitrite within the bacterial cellular compartments that are surrounded by membranes. The nitrite is then released from the bacterial cell and can be used by the foraminiferal host. The latter scenario may be more beneficial for the bacterium. Overall, our results suggest that the *Globobulimina*-associated Desulfobacteraceae are able to reduce NO _3-_ to NO _2-_, and thus contribute to the foraminiferal denitrification by performing the first reaction in the denitrification pathway for which we found no evidence in the foraminiferal transcriptomes.

## Discussion

Our results demonstrate that foraminifera are habitat to bacterial communities that may play a role in their ability to thrive in oxygen-depleted habitats. A recent study demonstrates the endosymbiotic contribution to denitrification within a ciliate host (56). However, previous studies of denitrification in foraminifera argued against the possibility of bacterial contribution to foraminiferal denitrification (8, 11). Notably, most species in the foraminiferal microbiota are considered strict anaerobes, hence, when exposed to oxygen the foraminifera may lose their associated microbiota. Metagenomics sequencing of samples frozen directly after sampling is thus an important source of information on the composition and function of the foraminiferal microbiome.

Our results indicate that Desulfobacteracea members of the foraminiferal microbiota can utilize the NO _3-_ storage accumulated by their host. Whether the NO _3-_ reduction by bacteria is beneficial to the foraminifera remains, however, an open question. The bacteria may use the NO _3-_ it for their own respiratory processes or to build up organic compounds via the assimilation of NO _3-_. For example, it has been suggested that a proportion of NO _3-_ taken up by the foraminiferal species *Ammonia beccari* was used for amino acid synthesis, probably by resident bacteria (21). Previously we speculated about a foraminiferal sulfite reductase homolog performing the conversion of NO _3-_ (11). Yet, considering the absence of foraminiferal homologs to NO _3-_ reductase and the presence of NapA in all sampled Desulfobacteraceae MAGs, it is tenable to hypothesize that the reduction of NO _3-_ to NO _2-_ in foraminifera is performed by resident bacteria.

An association between foraminifera and Desulfobacteracea has been previously reported. For example, the presence of a putative Deltaproteobacterium (Desulfobacteracea) was previously described for *Virgulinella fragilis* (20), a foraminiferal species that we here predict to be a denitrifying species (Figure 3B). Furthermore, foraminifera from the genera *Ammonia, Elphidium* and *Haynesina* that are not expected to denitrify were found to harbor members of Desulfobacterales bacteria (25). Hence, the interaction between foraminifera and Desulfobacteracea may involve alternative nutritional currencies. Previous studies refer to the role of Desulfobacteracea in sulphur cycling and carbon/nutrient acquisition (20, 21, 25). Especially the carbon fixation capabilities via Wood-Ljungdahl pathway would be highly beneficial for the heterotrophic lifestyle of foraminifera, similarly to other symbioses that involve fixed-carbon as a nutritional currency [e.g., as in deep-sea mussels (57) or sponges (49)].

The radiation of early foraminiferal species has been estimated to occur between 690 and 1,150 million years ago (mya), while the first primitive rotaliids appeared in the late Permian around 260 mya (58, 59). Most of the rotaliids superfamilies diverged in the mid to late Triassic between approximately 240 and 200 mya (29, 60, 61) and the majority of extant species diverged during the Miocene (23.8 -5.5 mya) (62). Our phylogenetic analyses suggest that clades I and III diversified from within clade II, while the LCA of clade I probably originated from within clade III (Figures 2 & 3). Although most species reported to denitrify were reported from the Peruvian OMZ and Sweden, representatives of corresponding genera are found all over the world (Suppl. Table S5). Therefore, most extant rotaliids likely diversified from a denitrifying LCA far back in time. The origin of foraminiferal denitrification within or after diversification from members of clade II may coincide with the rapid increase of fossil records at the onset of the late Cretaceous ∼100 mya (60, 61). Our results therefore indicate that eukaryotic denitrification by rotaliids emerged late in foraminiferal evolution, possibly during the worldwide Cretaceous Ocean Anoxic Events (63, 64).

Considering the high conservation of nitrate transporters (Figure 1) and the observation of nitrate storage in many divergent species (9), it is tenable to speculate that foraminifera had the mechanisms for nitrate import and storage long before they evolved the ability to denitrify. NO _3-_ transporters are ancient eukaryotic enzymes that play a role in the NO _3-_ assimilation machinery and they are encoded in genomes of photoautotrophs such as plants and saprotrophs like fungi (65, 66). In contrast to bacteria, foraminifera (as most heterotrophs) are unable to perform NO _3-_ assimilation on their own. Thus, NO _3-_ accumulated by the foraminiferal hosts could have fuelled bacterial NO _3-_ metabolism of associated bacteria in exchange for organic compounds. We propose that denitrification by symbiotic bacteria was indeed the ancestral state of denitrification in foraminifera, similarly to gromiids (26), which share a common evolutionary origin with foraminifera (67). The finding of bacterial-like denitrification genes in rotaliids furthermore suggests that foraminifera may have acquired those genes from bacteria (11). Thus, a rare gene acquisition from a foraminifera-associated, denitrifying bacterium could have been at the origin of foraminiferal (eukaryotic) denitrification. Notably, the rotaliids denitrification gene set is incomplete and it varies among the species sampled here (i.e., *N. auris, B*.*plicata* or even *Rosalina* sp.). Species reported to release N _2_O instead of N _2_ (9) further support our observation. Thus, the presence of a partial denitrification pathway in rotaliids as well as in their resident Desulfobacteracea may suggest that the acquisition of denitrification ability in foraminifera occurred in multiple stages and is not yet complete.

## Methods

### Foraminifera sampling

Samples were collected off the Peruvian coast in the 2017 Austral winter (R/V Meteor M137) as described by Glock et al. 2019 (10). Briefly, sediment samples were taken with a video-guided sediment multiple corer (MUC) containing 6 liners along a depth transect at 12°S. The top 1 to 3 cm of the sediment cores were sampled and immediately wet sieved with surface water through staked sieves with mesh size of 2000μm to 63μm to retrieve benthic foraminifera. The foraminifera were rinsed in sterile seawater obtained by filtering core-overlying seawater with a sterile bottle top filtration system (Durapore filter, 0.2 µm) and a vacuum pump within 40 minutes of MUC arrival on deck. Focal species were manually picked using a stereomicroscope. These were characterised morphologically according to the literature (68-71). Up to 160 individuals (classified into focal species) were pooled in one cryo vial (2ml, RNAse free) and flash frozen in liquid nitrogen.

### Microscopy

Individual foraminifers were dehydrated in a graded ethanol series (70 %, 80 %, 90 %, 96 % and two times 100 %; 15 min each), air-dried at 20 °C for 12 h in a desiccator and mounted on aluminium stubs (PLANO GmbH) using conductive and adhesive carbon pads (PLANO GmbH). Subsequently, the preparations were sputter-coated with a 10-nm-thick gold-palladium (80/20) layer using a high vacuum sputter coater Leica EM SCD500 (Leica Microsystems GmbH) and visualised with a Hitachi S-4800 field emission scanning electron microscope (Hitachi High-Technologies Corporation) at an acceleration voltage of 3 kV and an emission current of 10 mA applying a combination of the upper detector and the lower detector. The *B. costata* specimen was taken from previous samplings by the General Direction of Research in Oceanography and Climate Change IMARPE (71). The corresponding sample was air-dried at 35–38 °C, sputter-coated with gold and visualised with an FEI Inspect S50 scanning electron microscope (Thermo Fisher Scientific Inc.) at an acceleration voltage of 3 kV.

### Nucleic acid extraction, and sequencing

Genomic DNA and total RNA from different biological samples were extracted using either the DNeasy® Plant Mini kit (QIAGEN) for genomics (DNA) or simultaneously purified using InnuPREP DNA/RNA Mini Kit (analytikjena) for transcriptomics (RNA). Foraminiferal cells in each samples (ca. 160 individuals) were disrupted by pestle-crashing on ice after immersion of the containing cryo-vial in liquid nitrogen. Samples for genomics were treated with lysozyme (200μl of 10mg/ml TE) and proteinase K (1mg/100μl). Samples for transcriptomics samples were treated only with lysozyme (6 ul of 20mg/ml TE) before additional crashing and 2-minute incubation was performed. After disruption and initial lysis, manufacturer protocols for the respective nucleic acid extraction were followed with modifications to the elution volumes (2 × 50μl for DNA and 2 × 25μl for RNA). Libraries for genomics were produced after DNA fragmentation (Covaris target 400, intensity 5, duty cycle 5% cycles per burst 200, 55sec treatment time) using NEBNext® Ultra II DNA Library Prep Kit for Illumina®. Whereas, transcriptomic libraries were produced using NEBNext® Ultra RNA Library Prep Kit for Illumina® (NEB) with mRNA isolation performed with poly-A mRNA beads. All libraries were produced in duplicates from two different sets of pooled individuals for each species and were produced without protocol interruption. Before sequencing, each library was quantified with a Qubit® fluorometer (Invitrogen by Life TechnologiesTM) and qualified using a TapeStation (Agilent technology). The libraries were sequenced paired-end (2 × 150 bp) on an Illumina® HiSeq 4000 platform.

### Sequencing datasets of foraminifera

Sequencing resulted in 4.6 billion paired-end reads covering 1.4 tera bases in total. This includes transcriptome and metagenome datasets (BioProject accessions PRJNA494828 & PRJNA503328). Reads were quality-checked by FastQC ver. 0.11.5 (http://www.bioinformatics.babraham.ac.uk/projects/fastqc; Aug 2016). Filtering and trimming of reads was performed using Trimmomatic (72) ver. 0.36 (Parameters: ILLUMINACLIP:primers.fa:2:30:10 LEADING:5 TRAILING:5 SLIDINGWINDOW:4:5 MINLEN:21; the file ‘primers.fa’ contained adaptor and contaminant sequences provided by Trimmomatic and FastQC). Processed reads from transcriptomes of the two samples per species were assembled into transcript contigs using SPAdes (73) ver. 3.11.1 (“--rna” option). Protein sequences were translated from transcripts as open reading frames (ORFs) using TransDecoder (74) ver. v5.0.2 (LongOrfs; “-m 30” option). Final protein names consist of the contig ids followed by the sequence positions covered by the CDS and an indicator for the forwards (+) or reverse (-) strand. Transcript abundance of individual transcriptome datasets are referring to Transcripts Per Million (TPM) determined by the Trinity pipeline (74) 2.4.0 (Trinity script ‘align_and_estimate_abundance.pl’) via RSEM (75) ver. 1.2.30 and Bowtie (76) 2.1.0 using paired-end reads. Additional raw sequencing reads were obtained from the Marine Microbial Transcriptome Project (MMETSP; SRA accessions: *Rosalina* sp., SRR1296887; *Sorites* sp., SRR1296734; *Ammonia* sp., SRR1300434; *Elphidium margaritaceum*, SRR1300475) and processed as described above. We found that the assembly obtained for *Sorites* sp. contained a high proportion of sequences likely originating from an algal species (Probably *Symbiodinium* sp.), which was removed by applying a binning approach considering only contigs with GC-content ≤38%. Finally, we included data for *Globobulimina* spp. sampled in Sweden from our previous study (Transcriptome data: GloT15) (11) and proteins annotated on the genome assembly of *Reticulomyxa filosa* (NCBI accession: GCA_000512085.1).

### Species phylogenies and rooting

Transcriptome completeness and heterogeneity was determined by assessing genome completeness via Benchmarking Universal Single-Copy Orthologs (77) (BUSCO v3; lineage ‘eukaryota’) method. Orthologous proteins determined as Complete (or Duplicated) by the BUSCO analysis were merged into protein clusters to study phylogenetic relationships among the species. In case of duplicated BUSCOs in metatranscriptomes, one representative was inferred based on sequence similarity. Therefore, all orthologs retrieved for the same metatranscriptome were compared to all members of the BUSCO protein cluster by global pairwise alignments using needle (EMBOSS tools version 6.6.0) (78). The ortholog with the highest median sequence similarity over all comparisons was picked as the representative sequence. Multiple sequence alignments used in the current study were obtained using MAFFT (79) (Version: 7; parameter: ‘linsi’) and the phylogenetic trees were reconstructed using IQTREE (80) (Version: 1.5.5 or 1.6.9; default parameters; note that ModelFinder is enabled by default). Phylogenies were reconstructed for the individual BUSCO clusters. Due to their morphological differences from other foraminiferal groups, monothalamids like *R. filosa* have been previously used as an outgroup for phylogenetic studies of foraminifera (29). Since the choice of a distantly related outgroup may lead to erroneous rooted topology due to long branch attraction (81), we further tested the robustness of the root position independent from outgroup species. Root positions were determined using MAD (28) (Parameters: ‘-bsnn’). The support in each root splits is calculated as the proportion of gene trees where the root position was inferred as such (28). For the overall species trees, multigene phylogenies were reconstructed using IQTREE (parameter: ‘-spp’) considering all protein cluster alignments of either foraminifera or Rotaliida.

Reference 18S rRNA sequences used for the large-scale phylogenies (Suppl. Table S5) were obtained from NCBI, MMETSP, SILVA (82), the foram barcoding project (http://forambarcoding.unige.ch/) and the Planktonic Foraminifera Ribosomal Reference database (83) (PFR^2^). These sequences were used as database sequences to identify 18S sequences from Peruvian species transcriptomes based on BLAST (84) (version: 2.2.28+; options: ‘-task blastn -evalue 1e^-10^’) searches. Among the hits retrieved we searched for a single representative transcript contig per transcriptome assembly. The transcripts with the highest product of sequence length and sequencing depth (i.e., coverage given by SPAdes) were determined as representative sequences for most of the focal species. However, for *B. costata, V. inflata, G. pacifica* and *N. auris* different representative sequences were chosen by giving sequence information (i.e., longer sequences) a higher weight than sequencing depth. To obtain 18S phylogenies first the reference sequences were aligned with MAFFT and subsequently representatives 18S rRNA sequences of the Peruvian transcriptomes were added (MAFFT options: ‘--addfragments --adjustdirection’) to the alignment followed by the tree reconstruction using IQTREE.

### Identification of foraminiferal denitrification proteins

To identify homologs to enzymes in the denitrification pathway, we used a similar approach as previously established in Woehle et al. 2018 (11) using the corresponding protein database expanded by the protein sequences identified for *Globobulimina* (11). The search for denitrification enzymes homologs in the transcriptome assemblies was performed with BLASTP (parameter: ‘-max_target_seqs 1000000 e-value 1e-5’). Protein sequences of hits with query coverage ≥ 40% and sequence identity ≥ 20% were extracted to obtain a first set of homologs. We further applied a cutoff to discard lowly represented transcripts with TPM < 2 in at least one of the two replicates sequenced. With the resulting protein set, we reiterated searches in the non-redundant NCBI protein (NR; version May 2018) and the RefSeq 88 database. (85) using diamond v0.9.22 applying the ‘--more-sensitive’ option. First best hit sequences per query were obtained and clustered with CD-HIT 4.6 (86) (option: “-c 0.98”) to reduce sequence redundancy. All obtained protein sequences of a given enzyme were aligned with MAFFT and phylogenetic trees were reconstructed with IQTREE (parameters: ‘-bb 1000

-alrt 1000’). The trees were rooted using an outgroup, if available, or MAD.

### Metagenomics processing

For the visualisation of metagenomic composition, trimmed paired-end reads from Peruvian species samples and the *Globobulimina* ‘Ambient’ samples (11) were subsampled via BBMap (version 36.84; https://sourceforge.net/projects/bbmap/; ‘reformat.sh’ script; parameters: ‘samplereadstarget=10000000 addslash=t’). The resulting reads were mapped against the NR database with ac-diamond (87) and classified using MEGAN6 (88) (Version 6.15.0; options ‘-a2t prot_acc2tax-Nov2018X1.abin -a2eggnog acc2eggnog-Oct2016X.abin -a2seed acc2seed-May2015XX.abin’). MEGAN6 was used for assessment of metagenomics communities and visual representation (See Figure 4) that were classified up to the order rank level with different taxon sets (e.g., all nodes or only the bacterial subtree). Metagenome assemblies were obtained by combining all trimmed reads per focal species using MEGAHIT (89) (Version 1.1.3; default settings). Individual bacterial genomes were obtained using the binning approach implemented with MaxBin2 (90) (Version 2.2.4; parameter: ‘-min_contig_length 500’) using coverage information of the two samples per species of foraminifera. Binning statistics were assessed using CheckM (91) (Version 1.0.11); a threshold for completeness of at least 80% and contamination of maximum 20% was applied to classify bacterial bins as draft genomes. The protein sequences obtained via checkM were used for further classifications. The average nucleotide identity (ANI) was calculated using a perl script obtained from https://github.com/chjp.

For the taxonomic assignment of genome bins, we used the diamond tool to find first best hits for each protein of a genome bin against the NR database (option: ‘-k 10’; *e*-value ≤ 1e-10) and retrieved the corresponding taxonomic assignments (as we previously described in ref. 11). For each bin, identical taxonomic hierarchies were counted with the genus being the lowest rank considered, sorted accordingly and stepwise searched for the lowest taxonomic rank supporting >50% of protein bin hits starting with the most abundant taxonomy. ‘Environmental samples’ and ‘Cellular organisms’ were not considered. For the phylogenetic reconstruction of Desulfobacteracea we determined protein families as protein clusters using sequences of Desulfobacteracea bins associated with *Globobulimina* and additional Desulfobacteracea genomes downloaded from NCBI. First, we determined reciprocal best BLAST hit pairs (rBBH; parameter: ‘-evalue 1e-5’) between all the Desulfobacteracea protein-coding sequences (92). Then rBBH pairs were globally aligned with the needle tool and pairs with ≥30% identical amino acids were sorted into clusters using the Markov clustering algorithm (93) (MCL; version 12-135). The 33 resulting protein clusters that contained a single copy for each of the Desulfobacteraceae strains (i.e., universal single-copy clusters) were aligned with MAFFT. The resulting alignments were concatenated and a splits network was reconstructed with SplitsTree (94) (version 4.15.1). Prediction of protein function for bacterial genomes were obtained using the KEGG Automatic Annotation Server (95) (KAAS; BBH method via BLAST and the following species set: hsa, dme, ath, sce, pfa, eco, sty, hin, pae, nme, hpy, rpr, mlo, bsu, sau, lla, spn, cac, mge, mtu, ctr, bbu, syn, aae, mja, afu, pho, ape, geo, dvu, dat, dpr, dol, dal, dak, dps, drt, dba, dao, dbr). Global pairwise identities for NapA protein sequences were inferred with needle. Signal peptides were predicted using the SignalP-5.0 webservice for gram-negative bacteria (96).

### Data availability

Sequencing reads are deposited at the single read archive (SRA) accessions SRR8144071 to SRR8144090 and SRR7971179 to SRR7971198. The assemblies are available at the transcriptome sequencing archive (TSA; See Suppl. Table 1 for accessions) and as whole genome shotgun (WGS) projects (See Suppl. Table 7 for accessions). All other information on accessing data analysed in this study is included in the manuscript or in the supplemental information.

## Acknowledgments

We thank Devani Romero Picazo and Anne Kupczok for critical comments on the manuscript. We gratefully acknowledge the scientific party and crew of the R/V Meteor cruise M137 as well as Asmus Petersen and Matthias Türk for their support at sea. Samples from Peru were obtained according to Peruvian access and benefit sharing regulations. We thank the Nagoya officer of Kiel University -Dr. Scarlett Sett -for her support of our research. We further thank Natalia Bernabe Lopez for her assistance in collecting metadata for the 18S sequences. The micrograph of *B. costata* was possible due to the General Direction of Research in Oceanography and Climate Change (IMARPE). Genome sequencing was performed in the Centre for Genome Analysis Kiel that is funded by the German Research Foundation (DFG). The study was supported by the German Research Foundation (DFG) via the SFB 754 on Climate–Biogeochemistry Interactions in the Tropical Ocean, the cluster of excellence The Future Ocean, and the European Research Council (Grant No. 281357 awarded to TD). C.W. was supported by the Kiel Life Science (KLS) Young Scientist Programme and would like to thank the Max Planck-Genome-centre Cologne for their support in data analysis.

## Author contributions

C.W., A.-S.R., N.G., T.W., J.W., D.R., J.S. & T.D. designed the research strategy and performed sampling. C.W. analysed the transcriptomes and metagenomes. A.-S.R. performed the experimental lab work. C.H. constructed incubation chambers. D.R. produced the scanning electron micrograph of *B. costata*. J.M. produced all other scanning electron micrographs, which was supported by S.N.G. C.W. and T.D. wrote the manuscript with input from all co-authors.

## Supplementary material

**Suppl. Figure S1.**
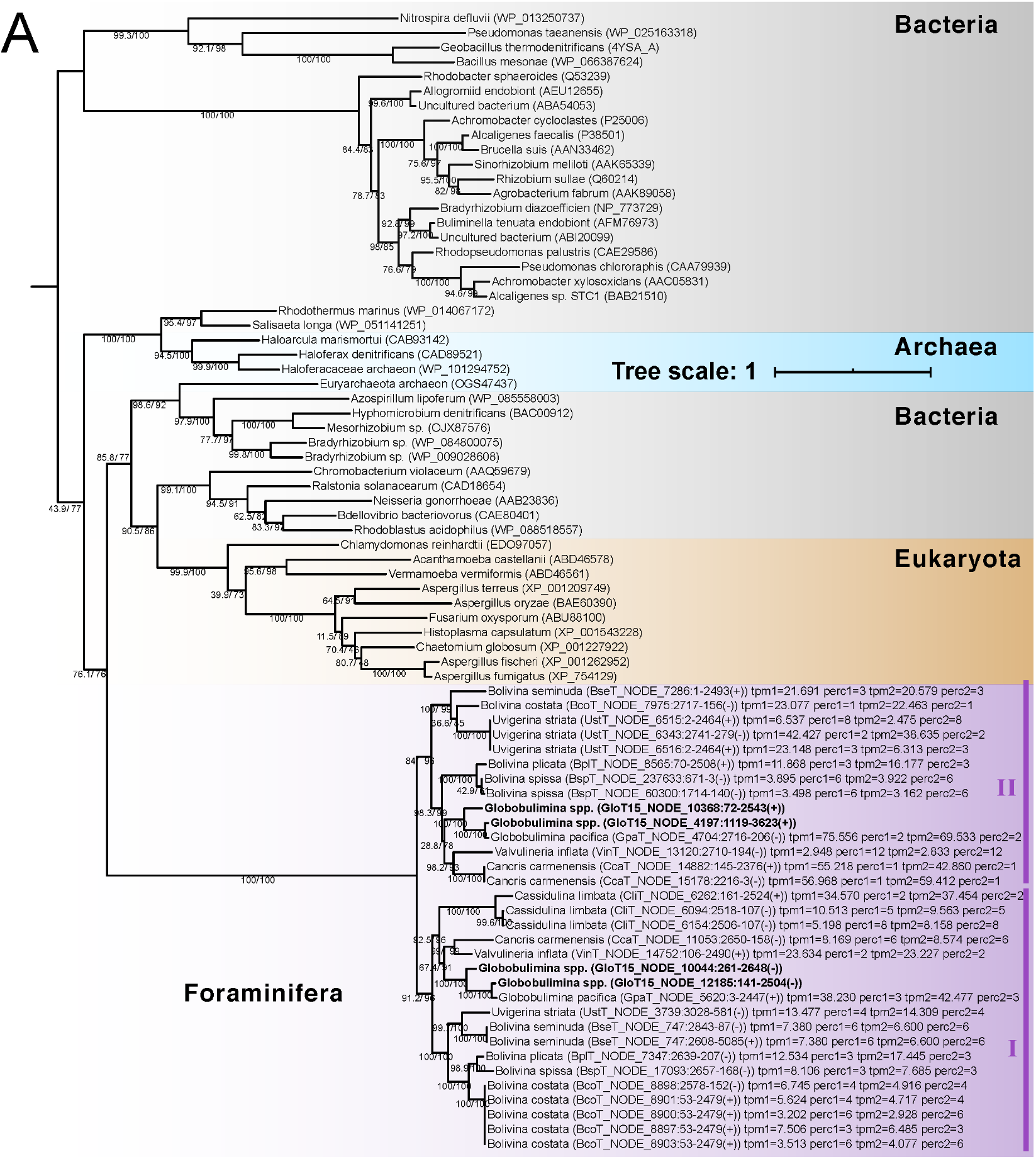
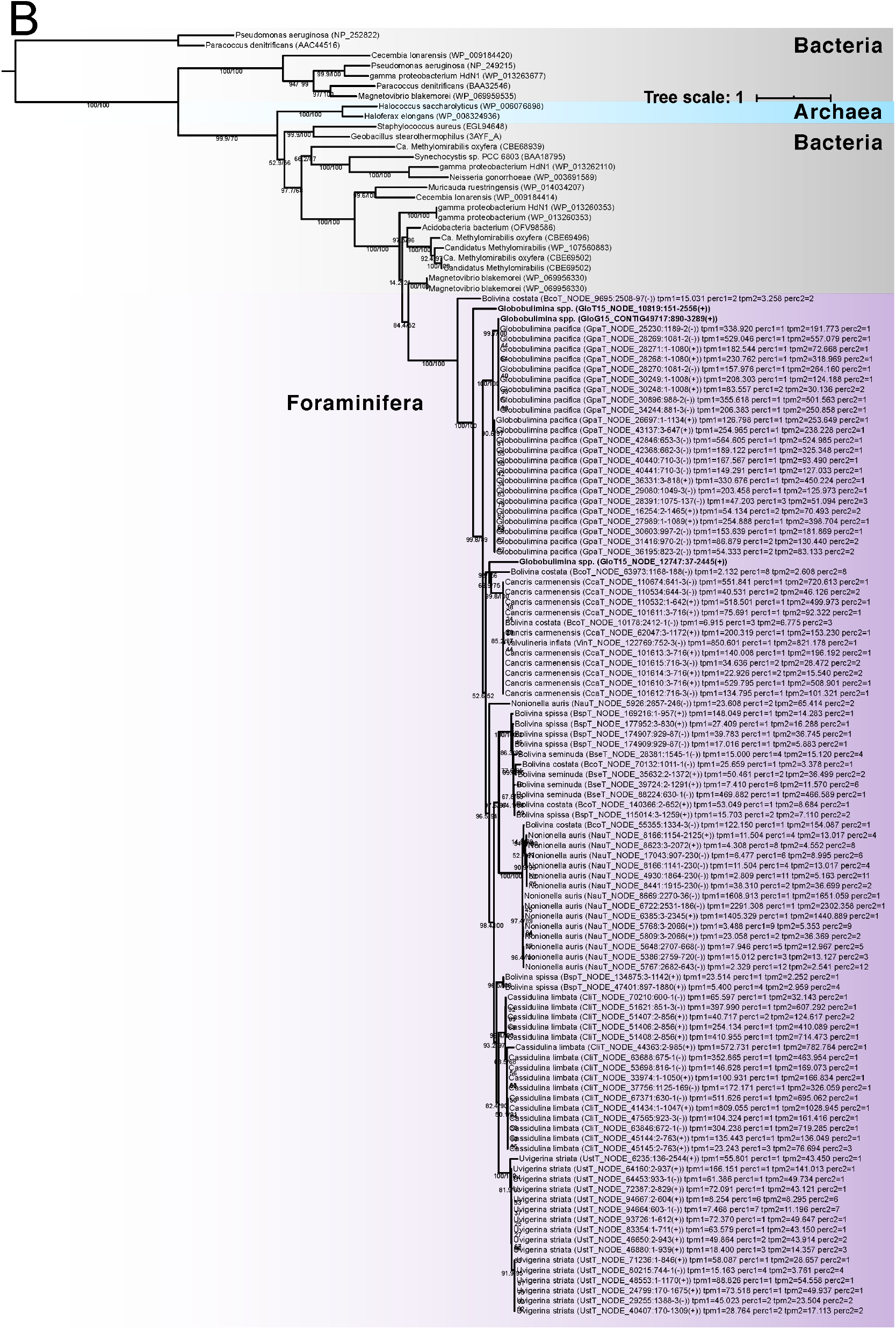
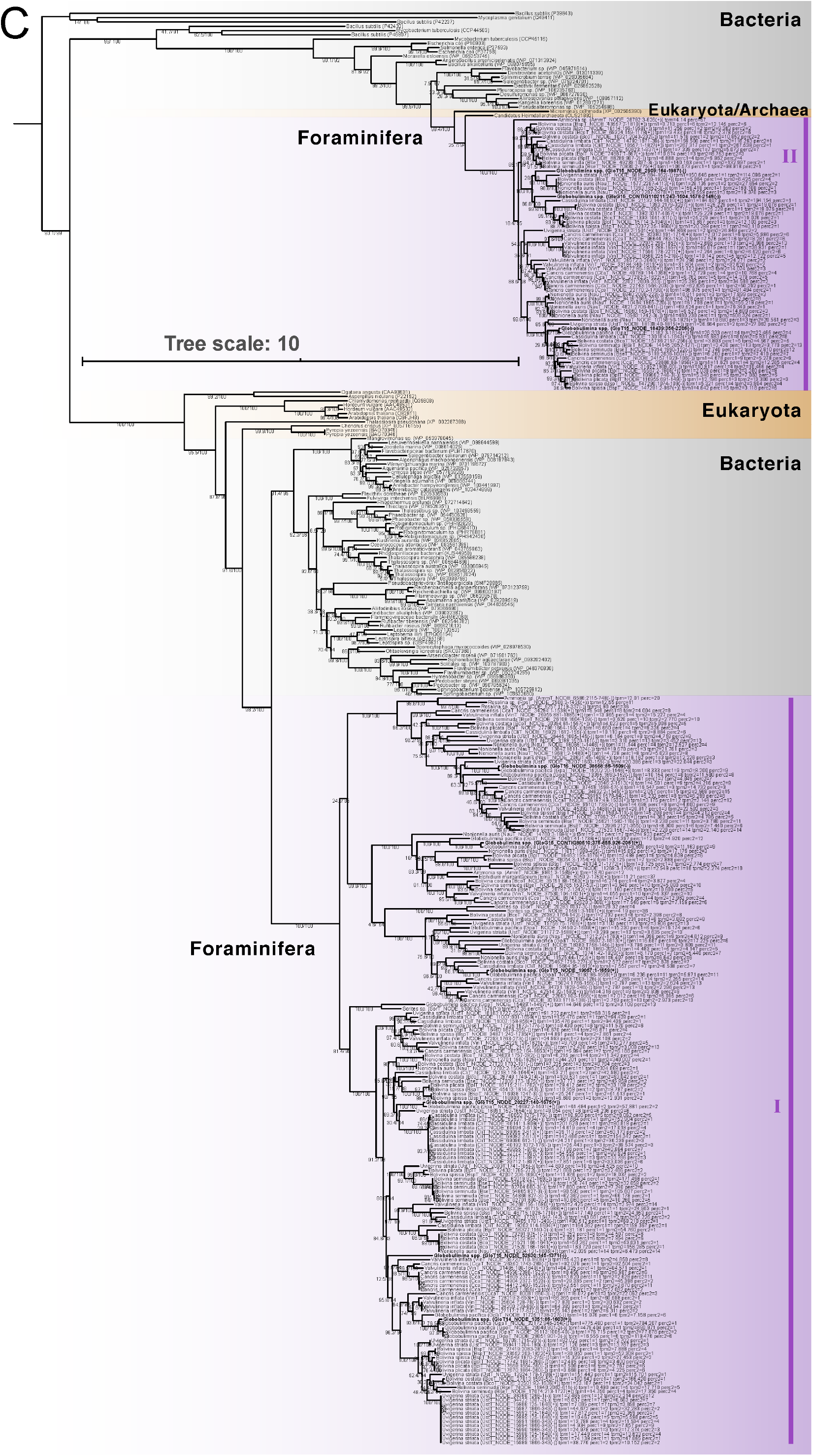
Scanning electron micrographs of sampled species. Detailed views of the graphs shown in Figure 1A.

**Suppl. Figure S2.**
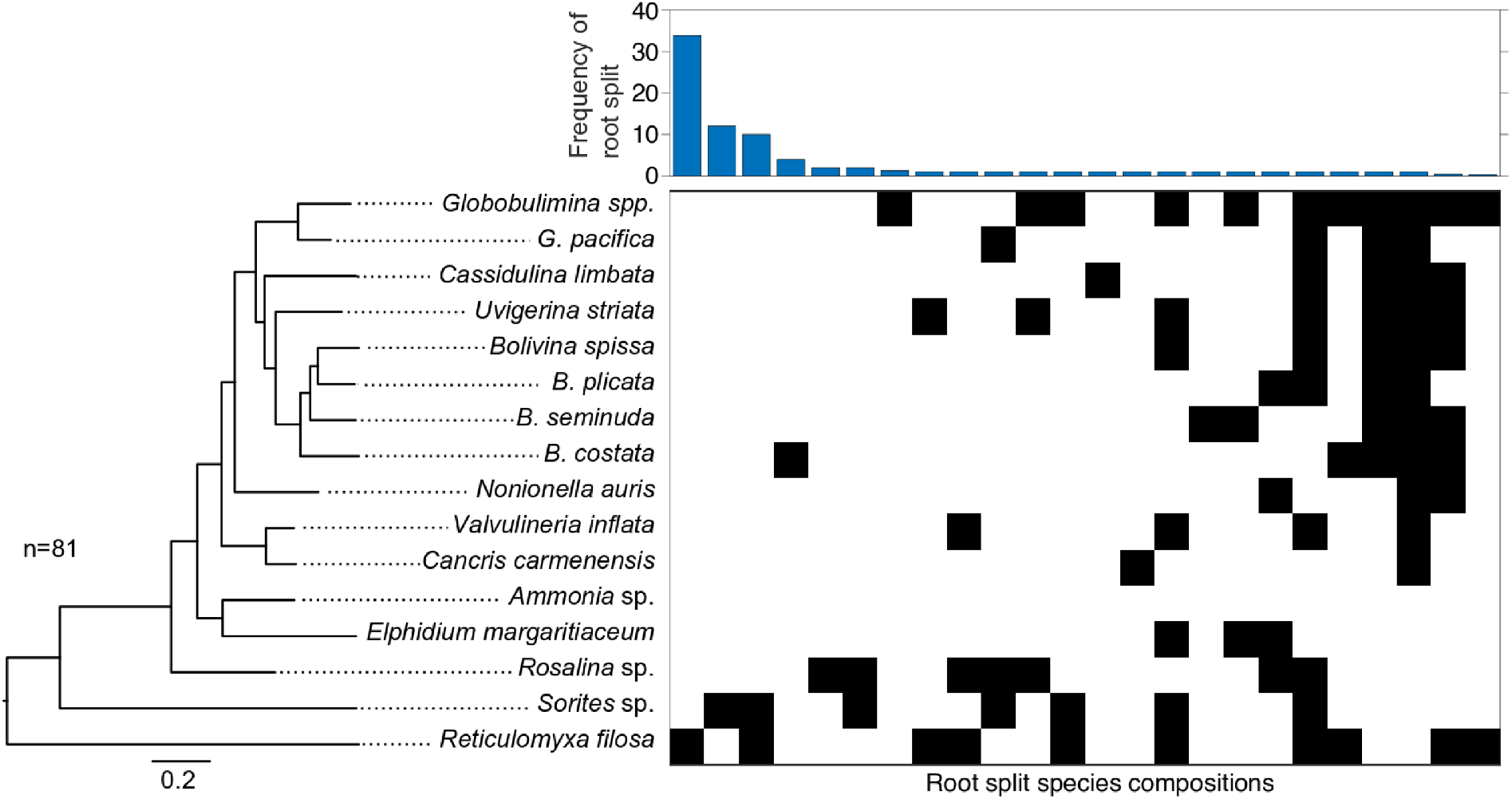
Complete phylogenies of NirK, Nor and Nrt homologs. Phylogenetics trees of the foraminiferal denitrification proteins A) NirK B) Nor and C) Nrt that survived the cutoff together with homologs from public databases. Transcriptome labels contain transcription values of the two replicates (tpm1 & tpm2) as well as their percentile expression rank within corresponding transcriptomes. The labels contain as well the transcript id, region and orientation for individual open reading frames used. Numbers of the branches represent statistical support by ultrafast bootstrap and SH-like approximate likelihood ratio test results (following the ‘/’) each with 1000 replicates. Branch support values of zero are omitted. If only one value is shown it represents the SH-like approximate likelihood ratio test. The background colour coding indicates different taxonomic groups. *Gobobulimina* spp. homologs from Woehle et al. 2018 are shown in bold. Further phylogenetic (sub-)clades, resembling those from Woehle et al. 2018, are highlighted by purple bars. The Nor tree was rooted using two cbb3 oxidases as outgroup, while for the Nrt tree representatives of DHA14 and ACS family of MFS transporters as outgroup. The NirK tree was rooted using the MAD method.

**Suppl. Figure S3.**
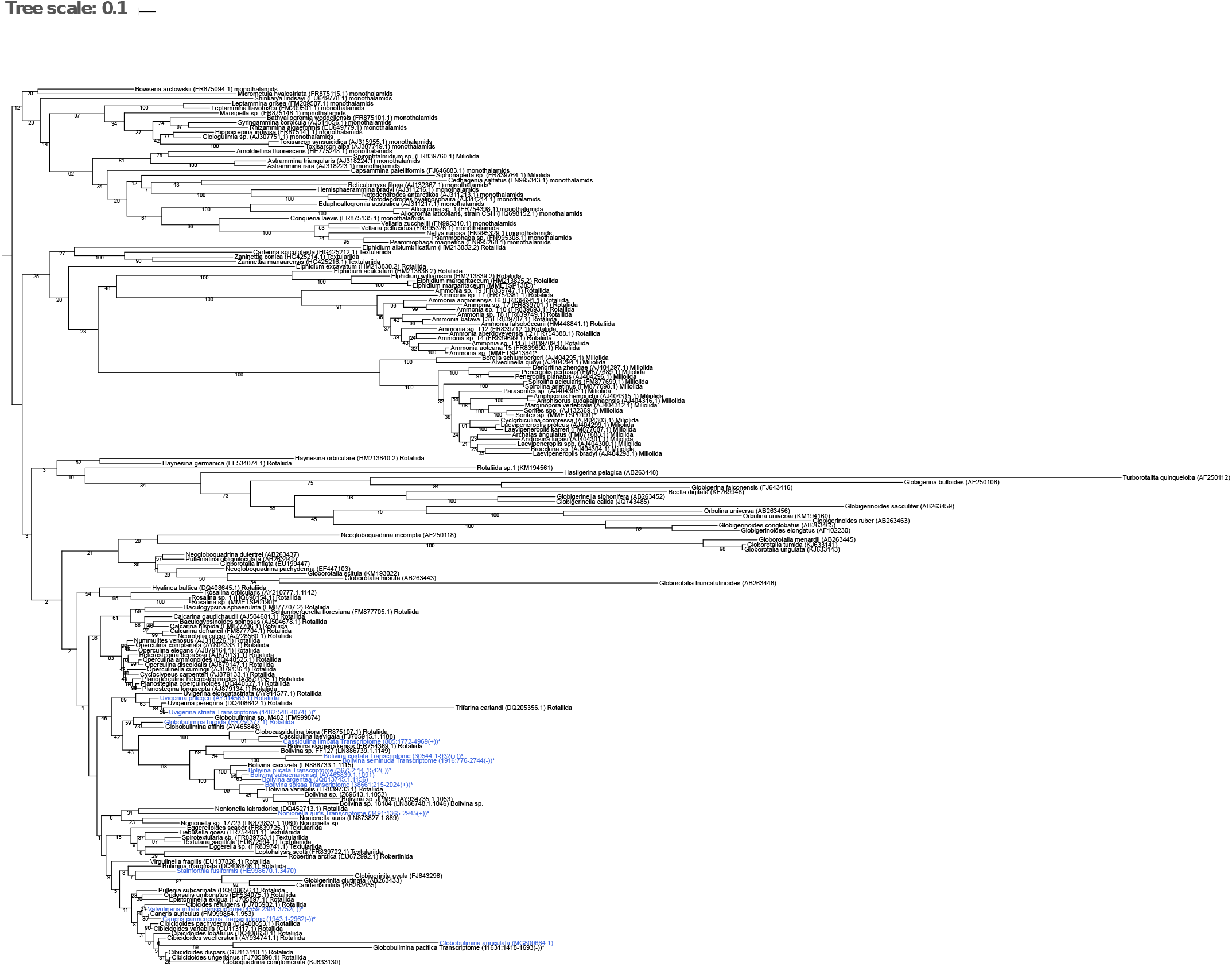
Foraminifera rooting based on single-protein trees. A Maximum-likelihood phylogenetic reconstruction of foraminifera species based on 81 eukaryotic protein marker sequences is shown on the left. Parametric bootstrap support is 1000/1000 at all the branches. Right of the phylogeny a split representation of root splits determined via the MAD approach for the 81 single gene trees is shown. Each column represents a root. Root branches are reported as separation of two groups (black and white boxes) indicating for the species found on either side of the root split. The bar graph (top) reports the single gene tree count supporting the corresponding root splits. These were ranked by frequency.

**Suppl. Figure S4.** Uncollapsed 18S phylogeny of foraminifera. Species shown to denitrify experimentally are highlighted in blue. The only exception is *Stainforthia fusiformis*, where denitrification activity has been shown for an unspecified species of the same genus. Species also shown in Figure 2 or Figure S1 are highlighted by asterisks. Bootstrap support values with 1000 replicates are shown at the branches. The trees were rooted by the clade of monothalamids containing *R. filosa*. The phylogeny corresponds to the one shown in Figure 3.

**Suppl. Table S1. Sequencing statistics for metagenomics & transcriptomics**.

**Supp. Table S2. List of foraminiferal homologs of denitrification proteins remaining after applying cutoffs**.

**Supp. Table S3. List of foraminiferal homologs of denitrification proteins removed by cutoffs**.

**Suppl. Table S4. List of marker proteins used for phylogenetic reconstructions. Suppl. Table S5. 18S sequences including metadata for phylogenetic reconstructions**.

**Suppl. Table S6. Proportions of family rank classification of foraminifera-associated microbial communities**.

**Suppl. Table S7. High quality genomic bins of foraminifera-associated species communities**.

**Suppl. Table S8. Metabolic classification of Desulfobacteracea genomes**.

